# Synthetic torpor triggers a neuroprotective and regulated mechanism in the rat brain, favoring the reversibility of Tau protein hyperphosphorylation

**DOI:** 10.1101/2022.03.25.485745

**Authors:** Fabio Squarcio, Timna Hitrec, Emiliana Piscitiello, Matteo Cerri, Catia Giovannini, Davide Martelli, Alessandra Occhinegro, Ludovico Taddei, Domenico Tupone, Roberto Amici, Marco Luppi

**Affiliations:** Department of Biomedical and NeuroMotor Sciences, University of Bologna, Bologna, Italy; Department of Neurological Surgery; Oregon Health & Science University; Portland, OR – USA; Department of Experimental, Diagnostic and Specialty Medicine, University of Bologna, Bologna, Italy; Centre for Applied Biomedical Research – CRBA, University of Bologna, St. Orsola Hospital, Bologna, Italy

**Keywords:** Deep hypothermia, microtubules, melatonin, GSK3β, hippocampus, parietal cortex

## Abstract

Hyperphosphorylated Tau protein (PPTau) is the hallmark of tauopathic neurodegeneration. During “synthetic torpor” (ST), a transient hypothermic state which can be induced in rats by the local pharmacological inhibition of the Raphe Pallidus, a reversible brain Tau hyperphosphorylation occurs. The aim of the present study was to elucidate the – as yet unknown – molecular mechanisms underlying this process, at both a cellular and systemic level. Different phosphorylated forms of Tau and the main cellular factors involved in Tau phospho-regulation were assessed by western blot in the parietal cortex and hippocampus of rats induced in ST, at either the hypothermic nadir or after the recovery of euthermia. Pro- and anti-apoptotic markers, as well as different systemic factors which are involved in natural torpor, were also assessed. Finally, the degree of microglia activation was determined through morphometry. Overall, the results show that ST triggers a regulated biochemical process which can counteract PPTau formation starting, unexpectedly even for a non-hibernator, from the hypothermic nadir. In particular, at the nadir, the glycogen synthase kinase-β was largely inhibited in both regions, the antiapoptotic factor AKT was significantly activated in the hippocampus, and melatonin plasma levels were significantly increased, while a transient neuroinflammation was observed during the recovery period. Together, the present data suggest that ST can trigger a previously undescribed latent and regulated physiological process, that is able to cope with brain PPTau formation.

**Graphical abstract:** 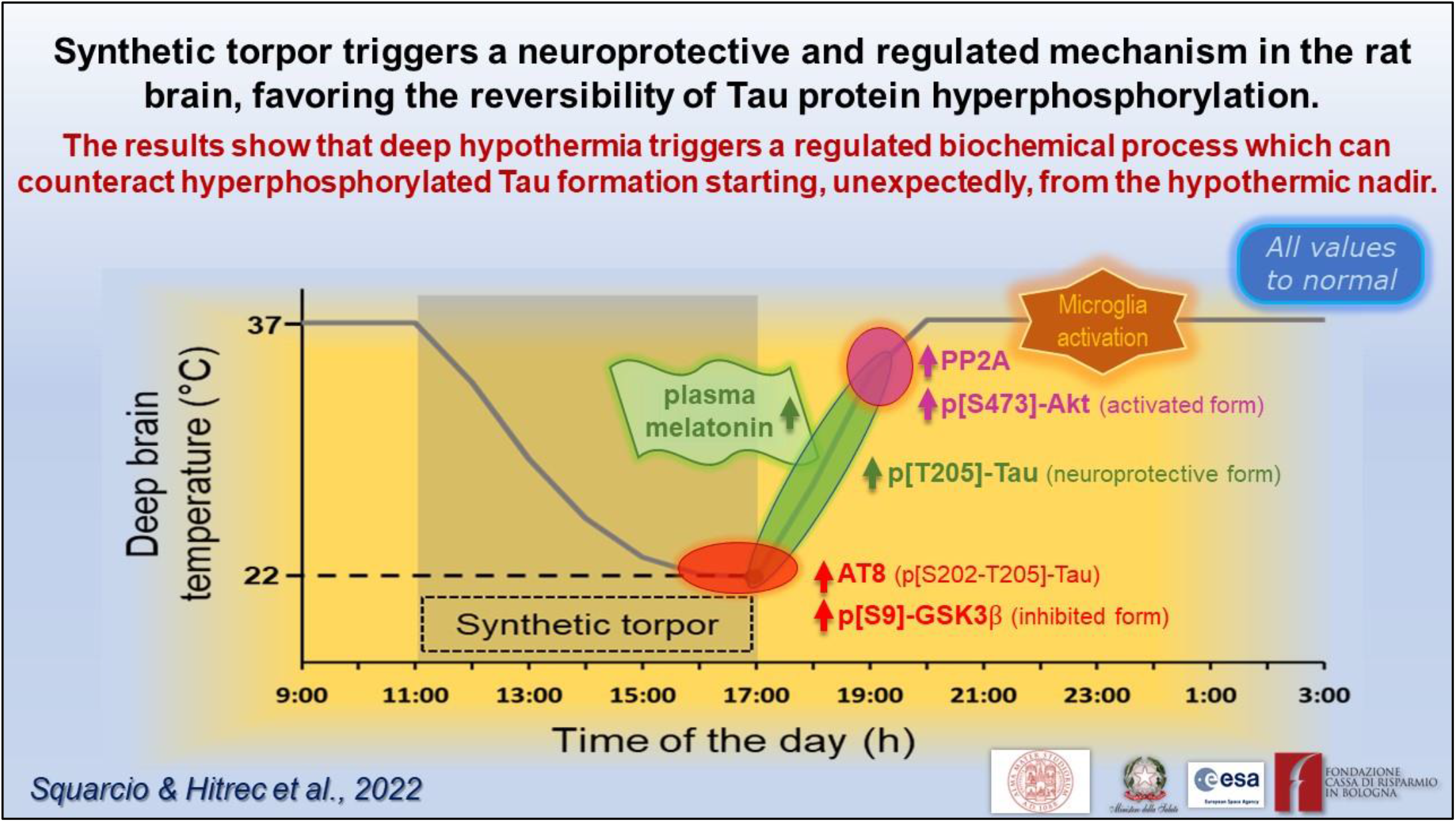

## Introduction

The Tau protein has a fundamental function in neurons (Wang and Mandelkow, 2016), the name itself deriving from the very first description of the key role it plays in the assembly and stabilization of microtubules (MTs) (Weingarten et al., 1975). When it is hyperphosphorylated, Tau (PPTau) loses its primary function: Tau monomers detach from MTs, showing a tendency to aggregate in oligomers and then evolve toward the formation of neurofibrillary tangles (Gerson et al., 2016; Wang and Mandelkow, 2016). This mechanism represents the main pathological marker of neurological diseases that are also termed as tauopathies (Kovacs, 2017), including Alzheimer’s disease (AD) and other neurodegenerative disorders (Wang and Mandelkow, 2016; Kovacs, 2017).

The formation of PPTau is not exclusive to neurodegenerative diseases. As a matter of fact, in response to hypothermic conditions, PPTau is also abundantly expressed in the brain. This is the case of hibernation (Arendt et al., 2015), deep anesthesia (Whittington et al., 2013), and “synthetic torpor” (ST; Luppi et al., 2019), a condition the rat, a non-hibernator, enters into in response to pharmacological inhibition of thermogenesis (Cerri et al., 2013; Cerri, 2017). In almost all these cases, with the only exclusion of anesthesia-induced hypothermia in transgenic mice models of tauopathy (Planel et al., 2009), Tau hyperphosphorylation is reversible and not apparently leading to neurodegeneration. Since mice are facultative heterotherms, able to enter torpor if necessary (Hudson and Scott, 1979; Oelkrug et al., 2011; Hitrec et al., 2019), the PPTau reversibility observed during ST in the non-hibernator (Luppi et al., 2019; Hitrec et al., 2021) appears of particular relevance from a translational point of view.

The physiological mechanisms responsible for Tau hyperphosphorylation at a low body temperature, and its return to normality in euthermic conditions, are not yet well understood. This process could be theoretically explained by simply considering the physical chemistry of the two main enzymes involved in Tau phospho-regulation: glycogen-synthase kinase 3β (GSK3β) and protein-phosphatase 2A (PP2A; Planel et al., 2004; Su et al., 2008). Lowering of the temperature induces the inactivation of both the enzymes, but with different kinetics: PP2A deactivates faster than GSK3β (Planel et al., 2004; Su et al., 2008). Thus, lowering of the temperature might allow GSK3β to act on Tau filaments without being counter-balanced by PP2A, at least for a while. However, this cannot explain the rather fast dephosphorylation of PPTau observed during recovery from ST (Luppi et al., 2019; Hitrec et al., 2021).

The aim of the present work was to verify that the hyperphosphorylation and subsequent dephosphorylation of Tau, during and after ST, come from active molecular mechanisms triggered by deep hypothermia, at both cellular and systemic levels, not only being the result of temperature-dependent enzymatic activity modifications. Therefore, levels of different cellular factors in the parietal cortex (P-Cx) and in the hippocampus (Hip) of animals in ST and in normothermic conditions were compared. In particular, we determined levels of: i) AT8 (p[S202/T205]-Tau [Malia et al., 2016]) and Tau-1 (non-phosphorylated Tau); ii) GSK3β (total and inhibited form); iii) Akt (total and activated form; also known as protein-kinase B); iv) PP2A. Moreover, since Tau can be phosphorylated at different residues (Wang and Mandelkow, 2016), the levels of p[T205]-Tau form, which has been shown to have a neuroprotective role (Ittner et al., 2016), was also determined. Neuroinflammation, an important feature of tauopathies (Ransohoff, 2016; Nilson et al., 2017), was also assessed by analyzing the degree of microglia activation, together with that of pro-apoptotic and anti-apoptotic factors. Furthermore, considering that melatonin and noradrenaline might have a neuroprotective role (Herrera-Arozamena et al., 2016; Shukla et al., 2017; Benarroch, 2018) and their release is strongly affected, respectively, in hibernators during arousal from torpid bouts (Stanton et al., 1986; Willis & Wilcox, 2014), and, in general, during a strong thermogenic activation (e.g., such as that needed when returning to euthermia [Braulke and Heldmaier, 2010; Cerri et al., 2013; Cerri et al., 2021]), we considered these systemic factors worth studying. Therefore, plasma levels of melatonin and noradrenaline, together with those of adrenaline, dopamine, cortisol, and corticosterone were also determined.

In this study we confirm that ST is associated with the hyperphosphorylation of Tau and that this process is reversed during the return to euthermia. Moreover, the return to normal phosphorylation levels of Tau is not merely due to temperature changes but is also associated with an active neuroprotective biochemical process involving the inhibition of GSK3β, the activation of Akt and an increase in plasma levels of melatonin. This biochemical process is triggered by deep hypothermia, since rapidly elicited at the lowest brain temperature reached during ST. Lastly, it results that neuroinflammation induced by ST is mild and transitory.

## Material and methods

### Animals

A total of 30 male Sprague–Dawley rats (250-350 gr; Charles River) were used. After their arrival, animals were housed in pairs in Plexiglas cages (Techniplast) under normal laboratory conditions: ambient temperature (Ta) set at 24 ± 0.5°C; 12h:12h light-dark (LD) cycle (L: 09:00 h–21:00 h; 100–150 lux at cage level); food and water *ad libitum*. All the experiments were conducted following approval by the National Health Authority (decree: No. 262/2020-PR), in accordance with the DL 26/2014 and the European Union Directive 2010/63/EU, and under the supervision of the Central Veterinary Service of the University of Bologna. All efforts were made to minimize the number of animals used and their pain and distress.

### Surgery

After one week of adaptation, rats underwent surgery as previously described (Cerri et al., 2013). Briefly, deeply anesthetized rats (Diazepam, 5 mg/kg i.m.; Ketamine-HCl, Imalgene 1000, Merial, 100 mg/kg, i.p.) were placed in a stereotaxic apparatus (David Kopf Instruments) and surgically implanted with: (i) electrodes for electroencephalogram (EEG); (ii) a thermistor (Thermometrics Corporation) mounted inside a stainless steel needle (21 gauge) and placed beside the left anterior hypothalamus to record the deep brain temperature (Tb); (iii) a microinjection guide cannula (C315G-SPC; Plastics One; internal cannula extension below guide: +3.5 mm), targeted to the brainstem region involved in thermogenic control, the Raphe Pallidus (RPa), at the following coordinates from lambda: on the midline, 3.0 mm posterior and 9.0 ventral to the dorsal surface of the cerebellum (Paxinos & Watson, 2007; Morrison and Nakamura, 2019). Since reports showed that the inhibition of RPa neurons causes vasodilation (Blessing & Nalivaiko, 2001), the increase of tail temperature subsequent to the injection of GABA-A agonist muscimol (1 mM) in RPa was used as proof of the correct positioning of the guide cannula.

After surgery, rats received 0.25 ml of an antibiotic solution (penicillin G and streptomycin-sulfate, i.m.), analgesics (Carprofen—Rimadyl, Pfizer—5 mg/kg, i.m.), and 20 ml/kg saline subcutaneously.

Animals were constantly monitored until they regained consciousness and then left to recover for at least 1 week under standard laboratory conditions. The animal’s pain, distress, or suffering symptoms were constantly evaluated using the Humane End Point (HEP) criteria. 24 hours prior to the experimental session, rats underwent an adaptation period in a cage positioned within a thermoregulated and sound-attenuated box, at low Ta (15°C).

### Synthetic Torpor

To induce ST, we used a consolidated protocol (Cerri et al., 2013; Luppi et al., 2019; Tinganelli et al., 2019). Briefly, a microinjecting cannula was inserted into the previously implanted guide cannula. Then, 100 nl of muscimol (1 mM) was injected once an hour, six consecutive times. Following the last injection, Tb reached values of around 22°C (Cerri et al., 2013). The control group, conversely, was injected with artificial cerebrospinal fluid (aCSF; EcoCyte Bioscience). During the whole experiment, EEG and Tb signals were recorded, after being opportunely amplified, filtered, and digitalized (Cerri et al., 2013). The EEG registration, in particular, was used since of great help in monitoring brain functioning during ST, as shown in Cerri et al. (2013), and to exactly check for the starting point of the recovery period following ST (see later).

### Experimental Procedure

Animals were randomly assigned to five different experimental groups and were sacrificed at different times following the injection of either muscimol or aCSF (first injection at 11.00 a.m.). The experimental groups were the following (Figure 1):

**Figure 1.**
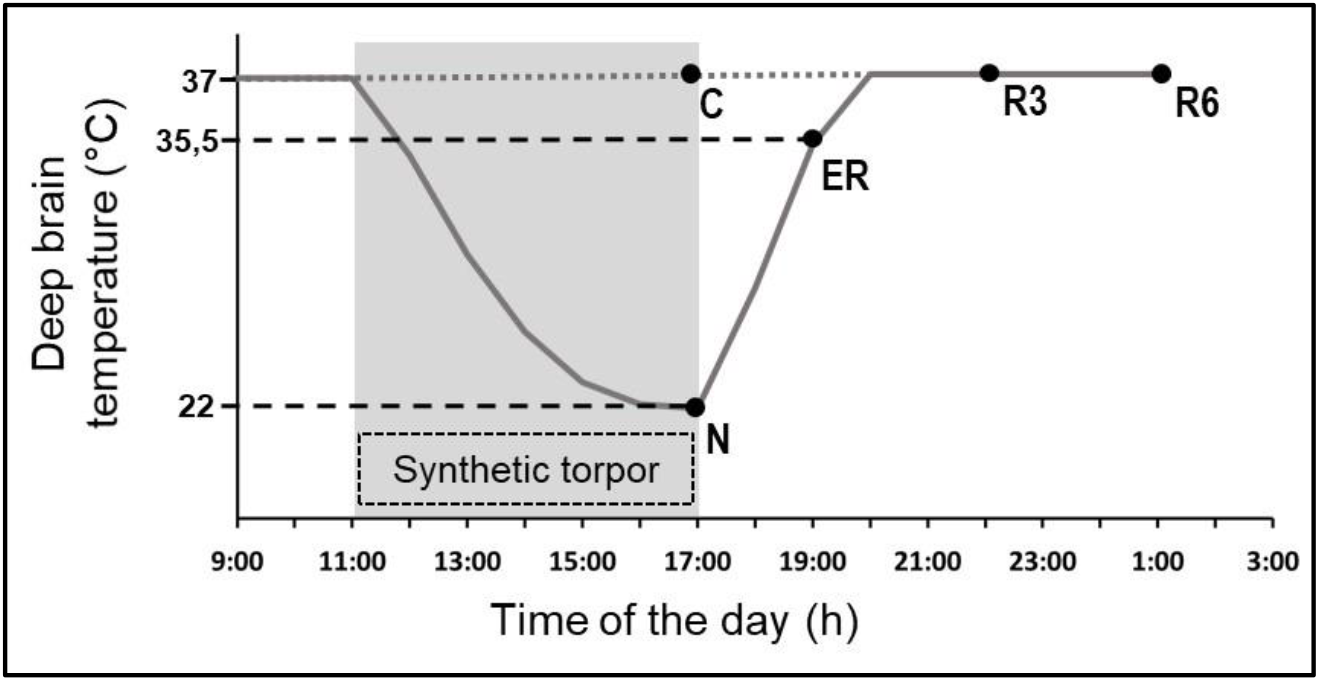
Schematic representation of the experimental conditions. The solid line indicates the progress of deep brain temperature (Tb) throughout the experiment. The dotted line refers to the control group (C; i.e., injected with artificial cerebrospinal fluid). The shaded area represents the period when synthetic torpor (ST) was induced (i.e., by injecting muscimol within the Raphe Pallidus). N, samples taken at nadir of hypothermia, during ST; ER, early recovery, samples taken when Tb reached 35.5 °C following ST; R3, samples taken 3h after ER; R6, samples taken 6h after ER. See Methods for details.

- C → Control, injected with aCSF and sacrificed at around 17.00 h, exactly matching the N condition (n = 6).
- N → Nadir, sacrificed 1 h after the last injection, at 17:00 h, when Tb reached the lowest temperature (i.e., the nadir) during hypothermia (n = 6; Tb = 22.8 ± 1.3 °C).
- ER → Early Recovery; sacrificed when Tb reached 35.5° C after ST, at around 19:00 h (n = 6).
- R3 → 3h Recovery, sacrificed 3 h after ER, at around 22:00 h (n = 6).
- R6 → 6h Recovery, sacrificed 6 h after ER, at around 1:00 h (n = 6).

The ER condition was empirically considered as the exact conclusion of the hypothermic period, since in this condition animals start to show normal EEG signals, including clear physiological signs of sleep (Cerri et al., 2013).

Blood was collected transcardially from each animal after induction of deep anesthesia, and was centrifuged at 3000xg for 15 min at 4°C; plasma was then separated for subsequent ELISAs. Three animals per condition were transcardially perfused with paraformaldehyde 4% (w/v) as previously described (Luppi et al., 2019; Hitrec et al., 2021), and brain excised for immunofluorescence (IF); three animals per condition were sacrificed by decapitation, and the fresh brain was extracted for subsequent western-blot (WB) analysis. All samples were stored at -80 °C until assayed.

### Immunofluorescence

The procedure has been described in detail in Luppi et al. (2019). Briefly, extracted fixed brains were post-fixed for 2 hours by immersion in the same solution used for the perfusion and put in a 30% (w/v) sucrose solution in phosphate buffer saline (PBS) and sodium-azide 0.02% (w/v) overnight for cryoprotection (Luppi et al., 2019). Thereafter, tissue samples were cut into 35 µm-thick coronal slices, using a cryostat-microtome (Frigocut 2800) set at -22.0 °C. All the slices were then stored at −80°C in a cryoprotectant solution: 30% (w/v) sucrose, 30% (v/v) ethylene glycol, 1% (w/v) polyvinylpyrrolidone in PBS.

Slices from approximately Bregma -2,0/-4,0 were used for free-floating immunostaining. These levels were chosen so that each slice would contain both a portion of the parietal cortex (P-Cx) and the hippocampus (Hip). Slices were rinsed twice in PBS and then incubated for 2 h in 1% (v/v) normal donkey serum. Slices were incubated overnight with polyclonal rabbit Anti-p[T205]-Tau (Thermo Fisher; 1:400) probed with a Donkey Anti-rabbit IgG conjugated with Alexa-594 (Thermo Fisher; 1:500).

Microglia activation was also assessed using the rabbit polyclonal Anti-Iba1 antibody (Wako Chemicals; 1:800) and the secondary antibody Donkey Anti-rabbit IgG conjugated with Alexa-594 (Thermo Fisher; 1:500), as previously described. Activation level was measured (ImageJ software, following appropriate calibration) using established morphometric parameters of microglia cells (Davis et al., 2017; Baldy et al., 2018): i) Soma area; ii) Arborization area; iii) Morphological index (MI): soma area/arborization area ratio; iv) Microglial density (counting the number of cells in every picture taken); v) Nearest neighbor distance (NND). This analysis was conducted on samples from the P-Cx and on only the CA3 field of Hip, since it represents the most vulnerable cortical field of the Hip in AD development (Padurariu et al., 2012; Ugolini et al., 2018).

Finally, endogenous levels of the large fragment (17/19 kDa) of activated Caspase-3, resulting from cleavage adjacent to Asp175, were detected using the rabbit polyclonal antibody for cleaved-Caspase 3 (Asp175) (Cell Signaling, 1:300), followed by a Donkey Anti-rabbit IgG conjugated with Alexa-594 (Thermo Fisher; 1:500).

Images were obtained with a Nikon eclipse 80i equipped with Nikon Digital Sight DS-Vi1 color camera, using 10x objective (20x for the microglia staining).

### Western Blots

After extraction, fresh brains were quickly transferred to a petri dish filled with ice cold PBS. P-Cx and Hip were then isolated, homogenized by ice-assisted sonication in lysis buffer condition (RIPA Buffer: 50 mM Tris buffer, 150 mM NaCl, 10% (v/v) NP-40, containing a cocktail of protease and phosphatase inhibitors [Sigma-Aldrich], 1 mM dithiothreitol [DTT] and 1 mM PMSF). The extract was centrifuged at 12000xg for 30 min at 4°C and stored at −80 °C. The protein concentration was determined using the Bio-Rad DC protein assay kit (Bio-Rad Laboratories). Aliquots were thawed on ice and then denatured in a sample buffer (containing: 500mM DTT, lithium dodecyl sulfate [LDS]) with Coomassie G250 and phenol red (Invitrogen™ NuPAGE) at 65 °C for 10 min. Then, protein samples (20 μg) were loaded and separated electrophoretically using a 1.0 mm thick 4 to 12 % Bis-Tris gel together with NuPAGE MOPS SDS Running Buffer (both by Invitrogen™ NuPAGE). The gels were then electrotransferred onto nitrocellulose membranes (Hybond C Extra, Amersham Pharmacia) via wet transfer. Membranes were blocked using 5 % (w/v) not-fat dry milk in 0,1% (v/v) tween-20 in PBS (PBST) for at least 40 min at room temperature, and then incubated overnight at 4 °C with the different primary antibodies indicated in Table 1. Bound antibodies were detected using horseradish peroxidase-conjugated Anti-rabbit and Anti-mouse secondary antibodies. The uniformity of sample loading was confirmed via Ponceau S staining and immunodetection of β-actin, used as a loading control. ChemiDoc™XRS+ (Image Lab™Software, Bio-Rad) was used to acquire digital images through a chemiluminescence reaction (ECL reagents, Amersham). A semi-quantitative measurement of the band intensity was performed using the same computer software and expressed as a ratio of band intensity with respect to the loading control, normalizing the different gels according to a randomly chosen sample used as an internal control (i.e., a sample taken from a single rat that was run on every gel for the different determinations).

**Table 1.** Primary antibodies employed in this study.

| Antibody | Type | Species | Specificity | MW (kDa) | Source and dilution |
| --- | --- | --- | --- | --- | --- |
| Anti-Total Tau | Mono- | M | several Tau protein isoforms between 50-70 kDa | 50-70 | Merck Millipore; WB 1:5000 |
| Anti-Tau-1 | Mono- | M | non phosphorylated 189-207 residues | 52-68 | Merck Millipore; WB 1:5000 |
| AT8 | Mono- | M | p-Tau (S202/T205) | 68-70 | Thermo Fisher; WB 1:1000 |
| Anti-p-Tau (T205) | Poly- | R | p-Tau (T205) | 68-70 | Thermo Fisher; IF 1:400; WB 1:1000 |
| Anti-GSK3β | Mono- | R | GSK3β | 46 | Cell Signaling Technology; WB 1:3000 |
| Anti-p-GSK3β | Mono- | R | p-GSK3β (S9) | 46 | Cell Signaling Technology; WB 1:3000 |
| Anti-Akt | Poly- | R | Total Akt1, Akt2, Akt3 | 56-60 | Cell Signaling; WB 1:2000 |
| Anti-p-Akt | Poly- | R | p-Akt1 (S473), p-Akt2 (S473), p-Akt3 (S473) | 56-60 | Cell Signaling; WB 1:2000 |
| Anti-GRP78 | Mono- | R | GRP78 | 78 | Cell Signaling; WB P-Cx 1:5000; WB Hip 1:3000 |
| Anti-XIAP | Mono- | M | XIAP | 54 | Santa Cruz Biotechnology; WB 1:1000 |
| Anti-PP2A Catalytic $\alpha$ | Mono- | M | PP2A | 36 | BDT Transduction Laboratories<sup>TM</sup>; WB 1:5000 |
| Anti-cleaved-Caspase 3 | Poly- | R | Cleaved Caspase 3 (A175) | 17-19 | Cell Signaling; IF 1:300; WB P-Cx 1:250; WB Hip 1:500 |
| Anti-β-actin | Mono- | M | β-actin | 42 | Thermo Fisher; WB 1:5000 |

Abbreviations: Mono-, monoclonal; Poly-, polyclonal; M, mouse; R, rabbit; MW, molecular weight; p-, phosphorylated; IF, Immunofluorescence; WB, Western Blot; P-Cx, Parietal Cortex; Hip, Hippocampus.

### ELISA determinations

Plasma samples from six animals for each experimental condition were pooled in order to obtain a sufficient volume to be assessed and with the aim to reduce the individual variability due to the relatively small number of animals. Therefore, we obtained two “grand-samples”, each assayed twice. Commercially available ELISA kits were used to measure plasma levels of melatonin (IBL International, RE54021), dopamine (IBL International, RE59161), adrenaline/noradrenaline (IBL International, RE59242), cortisol (IBL International RE52061), and corticosterone (Abnova, KA0468). All procedures were run following the manufacturer’s recommendations. Optical density was read with a spectrophotometer (Spark® microplate reader, Tecan).

### Statistical analysis

Statistical analysis was carried out using SPSS software (25.0). Western blot and ELISA results were analyzed using the Mann-Whitney non-parametric test, comparing every experimental condition with the relative C level. Microglia morphometric results were tested with a one-way ANOVA. In the case that ANOVA was significant, means of the different experimental conditions were compared with the relative C level, using the modified t-test (t*).

Immunofluorescence results were not statistically analyzed or considered as qualitative.

## Results

### Tau levels in the brain

The induction of ST did not induce changes in total Tau levels (Fig. 2-A), while AT8 (p[S202/T205]-Tau) and p[T205]-Tau showed a significantly higher peak in both P-Cx and Hip (P<0,05; for all comparisons), compared to C (Fig. 2, panels B and D). Interestingly, Figure 3 shows that the high level of p[T205]-Tau found in the Hip was specifically limited to the CA3 field (cf. Paxinos & Watson, 2007), not involving CA1 and CA2 fields. Moreover, during this condition Tau-1 (i.e., the non-phosphorylated form of Tau protein; Tab.1) mirrored the trend described for the phosphorylated forms, being significantly lower in both brain structures (P<0,05; for all comparisons) compared to control levels, as shown in Figure 2-C.

**Figure 2.**
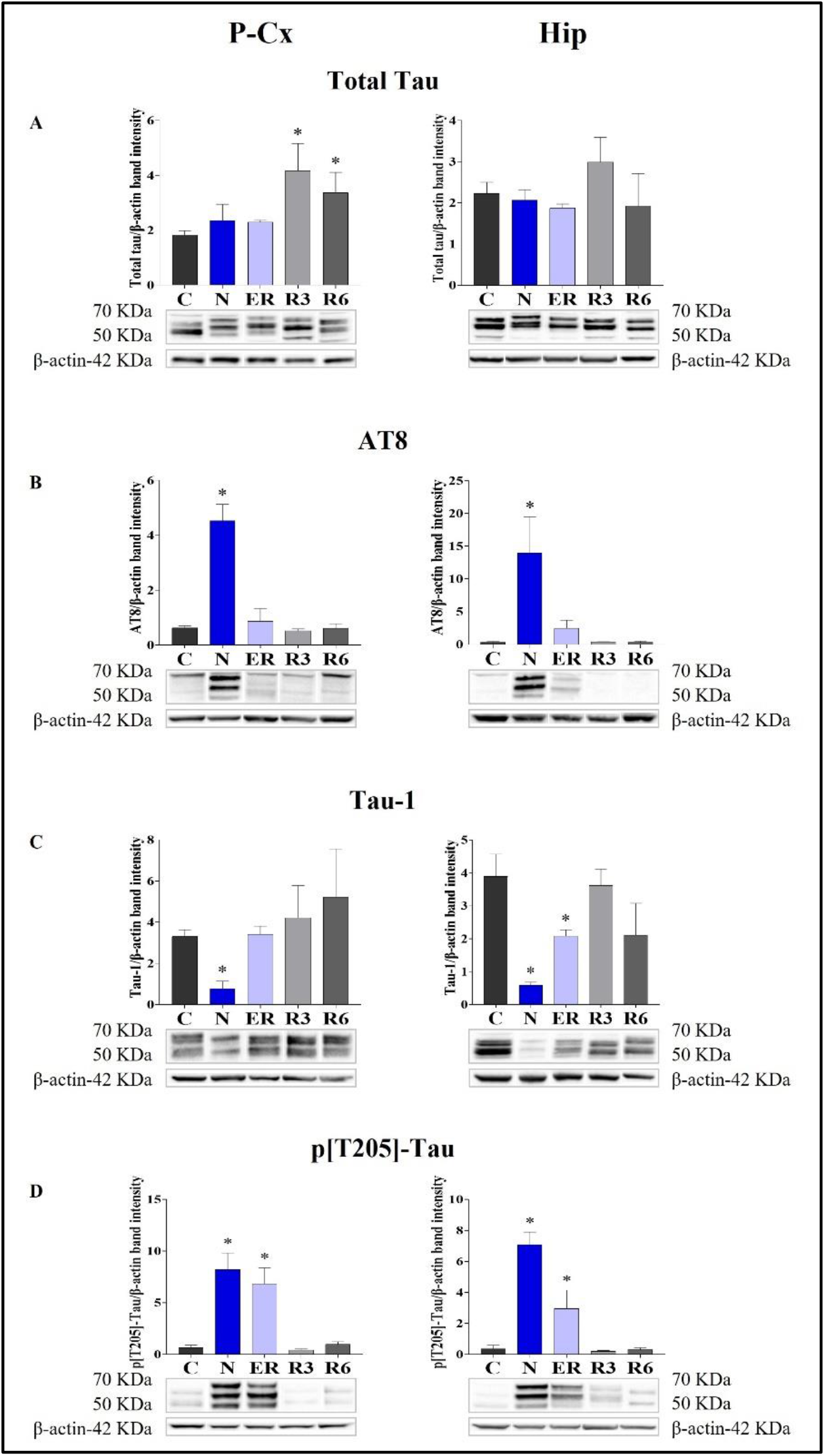
Western Blot detection of Total Tau and dephosphorylated/phosphorylated forms of Tau protein levels in brain extracts of the parietal cortex (P-Cx) and hippocampus (Hip). Below each histogram, WB representative samples are shown for each experimental condition. Panel A: Total Tau protein; Panel B: AT8 (Tau phosphorylated at Ser202 and Thr205); Panel C: Tau-1 (Tau dephosphorylated between 189 and 207 residues); Panel D: p[T205]-Tau (Tau phosphorylated at Thr205). Data are normalized by β-actin and expressed as means ± S.E.M., n = 3. *: p < 0.05 vs. C. Experimental groups (see Figure 1): C, control; N, samples taken at nadir of hypothermia, during ST; ER, early recovery, samples taken when Tb reached 35.5°C following ST; R3, samples taken 3h after ER; R6, samples taken 6h after ER.

**Figure 3.**
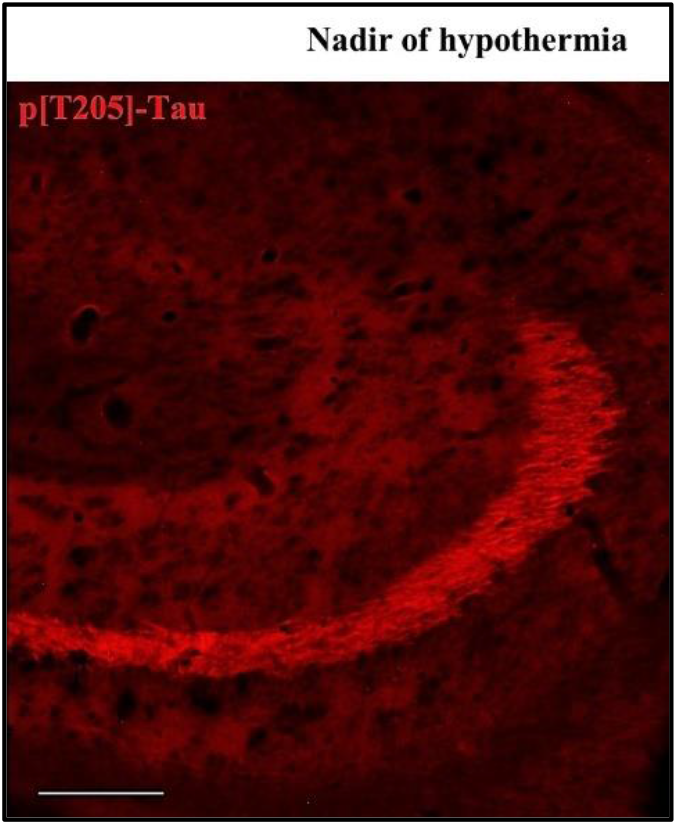
Representative pictures showing the CA3 hippocampus area stained for p[T205]-Tau, phosphorylated Tau protein in Thr-205 residue (secondary antibody conjugated with Alexa-594). The picture refers to the N condition (samples taken at the nadir of hypothermia, during the induction of synthetic torpor; see Figure 1). The picture represents the left hippocampal hemicortex, at Bregma -3,5. Calibration bar: 100 μm.

During the early stage of recovery (ER) AT8 returned to normal conditions (Fig. 2-B), while p[T205]-Tau level remained significantly higher than C (see Fig. 2-D) in both the brain structures studied (P<0,05; for all comparisons). Tau-1 levels returned to a normal condition in the P-Cx but was still significantly lower than control values in the Hip (P<0,05; Fig. 2-C). During the rest of the recovery period (R3 and R6) all the values affected by ST returned to normal levels (Fig. 2, panels B, C and D). In these recovery conditions, and only for P-Cx, total-Tau expression was found to be significantly higher than C (P<0,05; for both comparisons).

### Levels of kinases and phosphatases in the brain

Deep hypothermia induced different effects on GSK3β levels (Fig.4-A), depending on the brain structure considered. In fact, the expression of GSK3β was significantly lower in the P-Cx (N *vs*. C, P<0,05), but only showed an increasing trend in the Hip, that reached a significant level in ER (P<0,05). Consistently with what had been observed during ST, and in contrast with Hip, in P-Cx at ER the level of GSK3β was still lower than in C (P<0,05). Differently, results regarding the inhibited form of GSK3β (p[S9]-GSK3β; Fig. 4-B) showed a change, with a similar pattern in both the brain structures studied: values were significantly higher at N (P<0,05; for both P-Cx and Hip) compared to C, slowly returning to normal conditions during the recovery period (Fig. 4-B).

**Figure 4.**
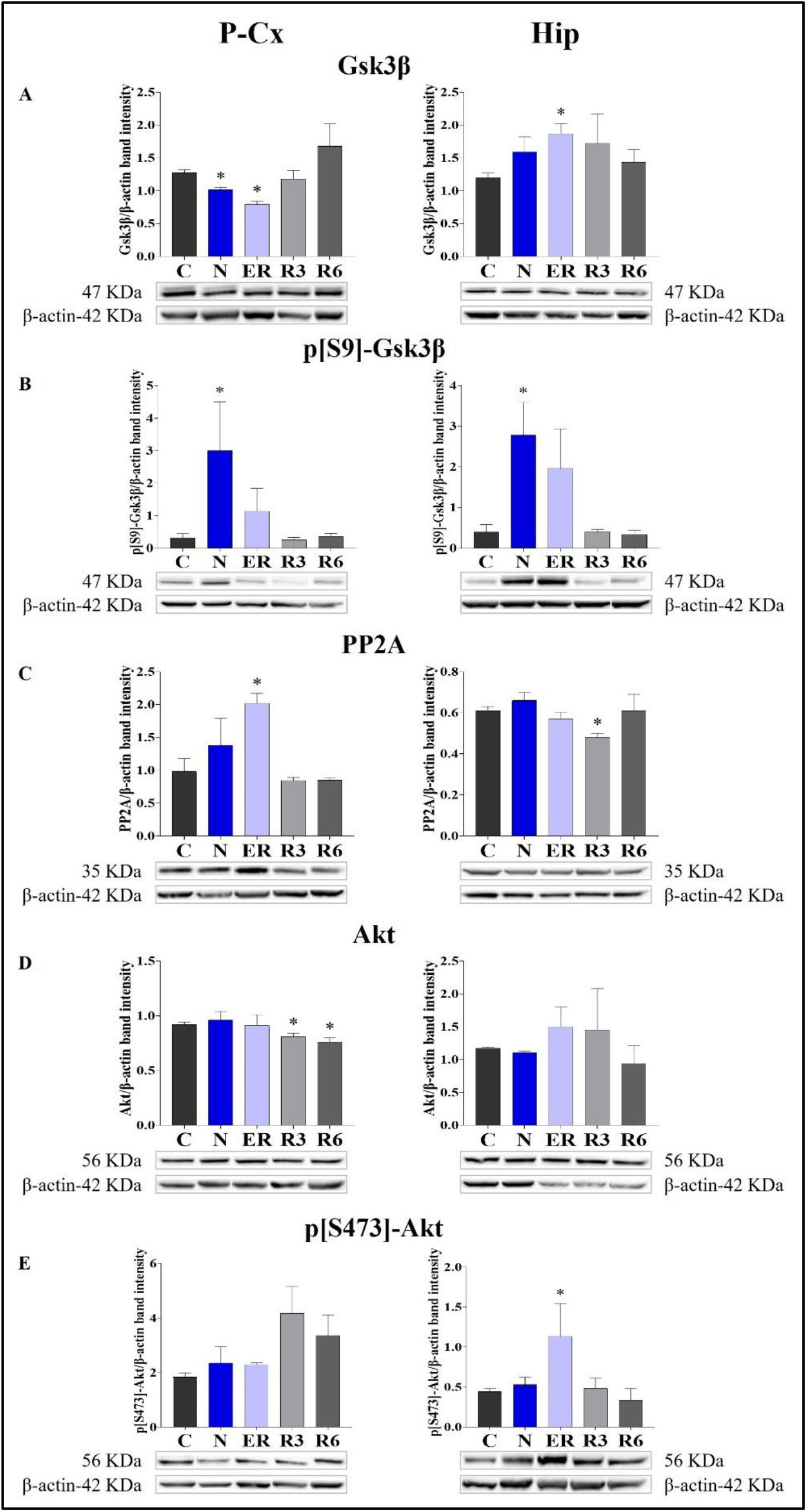
Western Blot detection of the main enzymes involved in phosphorylation and dephosphorylation of Tau, determined in brain extracts of the parietal cortex (P-Cx) and hippocampus (Hip). Below each histogram, WB representative samples are shown for each experimental condition. Panel A: glycogen-synthase kinase-3β (GSK3β), the main kinase targeting Tau; Panel B: p[S9]-GSK3β (inactive form of GSK3β, phosphorylated at Ser9); Panel C: protein phosphatase-2A (PP2A), the main phosphatase targeting Tau; Panel D: different isoforms of Akt (protein kinase-B; Akt 1/2/3), kinases targeting GSK3β at Ser9 and antiapoptotic factors; Panel E: p[S473]-Akt, the active form of Akt 1/2/3, phosphorylated at Ser473. Data are normalized by β-actin and expressed as means ± S.E.M., n = 3. *: p < 0.05 vs. C. Experimental groups (see Figure 1): C, control; N, samples taken at nadir of hypothermia, during ST; ER, early recovery, samples taken when Tb reached 35.5°C following ST; R3, samples taken 3h after ER; R6, samples taken 6h after ER.

As expected, PP2A (i.e., the main phosphatase acting on Tau protein) was significantly higher than in C (P<0,05; Fig. 4-C) during ER in the P-Cx, but such a trend was not observed in the Hip, where PP2A was lower than in C in R3 (P<0,05; Fig. 4-C).

Panels D and E within Figure 4, show total and active forms of Akt, respectively, also known as protein-kinase B, that play a role in neuroprotection and in contrasting apoptosis (Risso et al., 2015; Levenga et al., 2017). Total Akt expression was not affected by ST, and only at R3 and R6 within the P-Cx (Fig. 4-D) were levels significantly lower than in C (P<0,05; for both comparisons). The activated form of Akt (p[S473]-Akt) was induced only during the early stages of the recovery period (P<0,05; Fig. 4-E).

### Stressful/protective cellular markers in the brain

Taken together, all the results collected within Figure 5 show whether ST was stressful or protective at a cellular level. In particular: i) cleaved-Caspase 3 (panel A) is a factor that is commonly considered to be involved in the initiation of apoptosis (Fricker et al., 2018); ii) GRP78 (Glucose regulating protein 78; panel B) is a key factor that regulates the “unfolded protein response”, a well-recognized mechanism involved in cellular stress conditions (Ibrahim et al., 2019); iii) XIAP (X chromosome-linked inhibitor of apoptosis; panel C) is a factor that inhibits apoptosis (Holcik & Korneluk, 2001).

**Figure 5.**
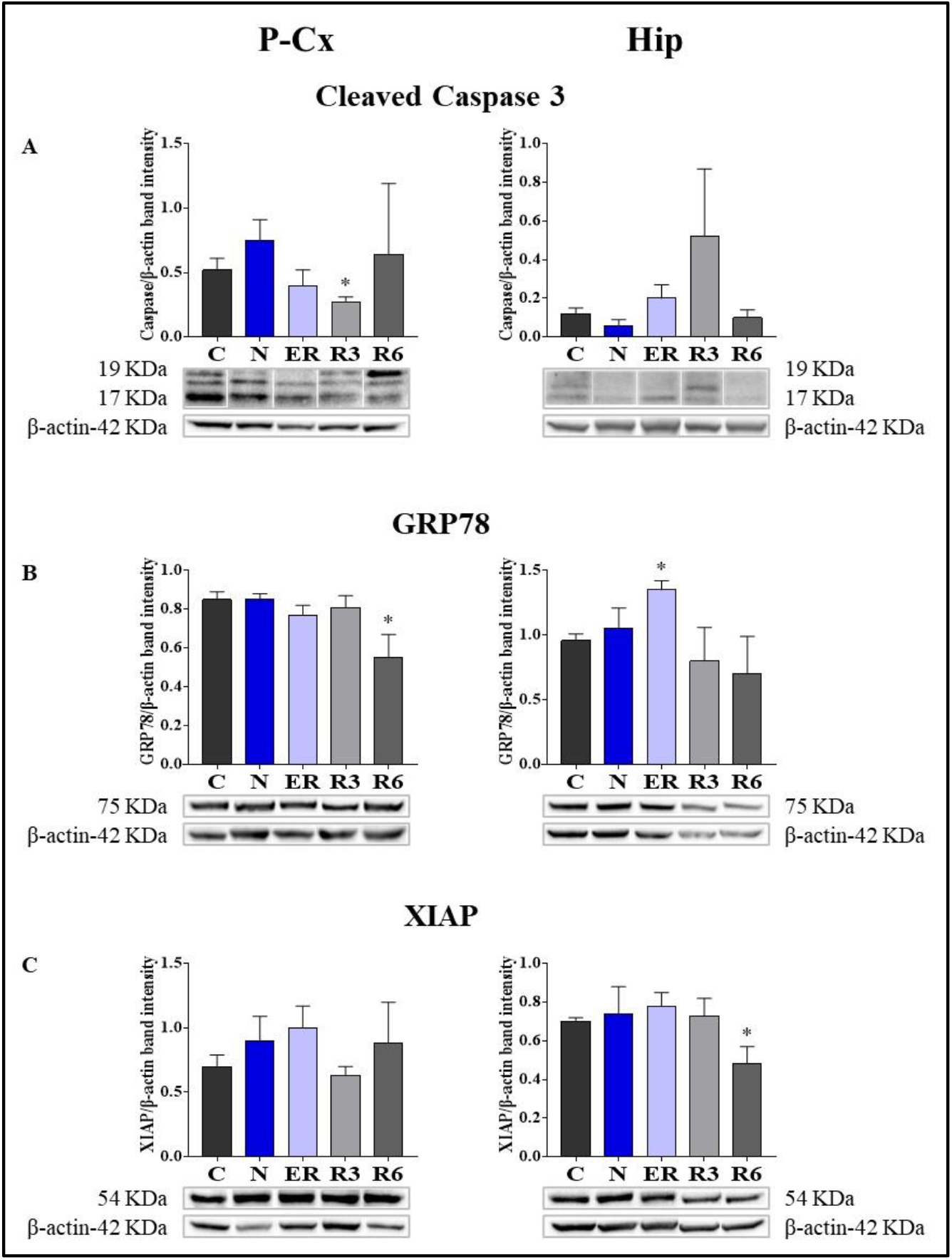
Western Blot detection of pro- and anti-apoptotic and cell-stress factors in brain extracts of the parietal cortex (P-Cx) and hippocampus (Hip). Below each histogram, WB representative samples are shown for each experimental condition. Panel A: cleaved-Caspase 3 (cleaved at Asp175 residue; i.e., the activated form); Panel B: Glucose regulating protein 78 (GRP78); Panel C: X chromosome-linked inhibitor of apoptosis (XIAP). Data are normalized by β-actin and expressed as means ± S.E.M., n = 3. *: p < 0.05 vs. C. Experimental groups (see Figure 1): C, control; N, samples taken at nadir of hypothermia, during ST; ER, early recovery, samples taken when Tb reached 35.5°C following ST; R3, samples taken 3h after ER; R6, samples taken 6h after ER.

These results show that the activated form of Caspase 3 (Fig. 5-A) was not affected by ST, resulting significantly lower (P<0,05) only in R3 within the P-Cx. GRP78 levels were found to be higher in ER within the Hip (P<0,05), but promptly returned to the normal condition during the rest of the recovery period. In P-Cx, this cellular factor presented significantly lower values in R6 (P<0,05), but maintained almost constant values throughout the other experimental conditions.

Finally, ST did not notably affect the neuroprotective factor XIAP. In this case, the only significant difference found was in R6 (P<0,05) within the Hip (Fig. 5-C), where it was lower than in C; while in the P-Cx XIAP levels were similar across the whole experiment.

### Systemic factors

Figure 6 shows the results obtained from plasma determinations of the systemic factors measured. While most of the results did not show significant variations across the experiment, melatonin and cortisol did. In particular, melatonin (Fig.5-A) was significantly higher in both N and ER, compared to C (P<0,05; for both comparisons), while plasma cortisol (Fig.5-E) was significantly lower than in C at ER (P<0,05).

**Figure 6.**
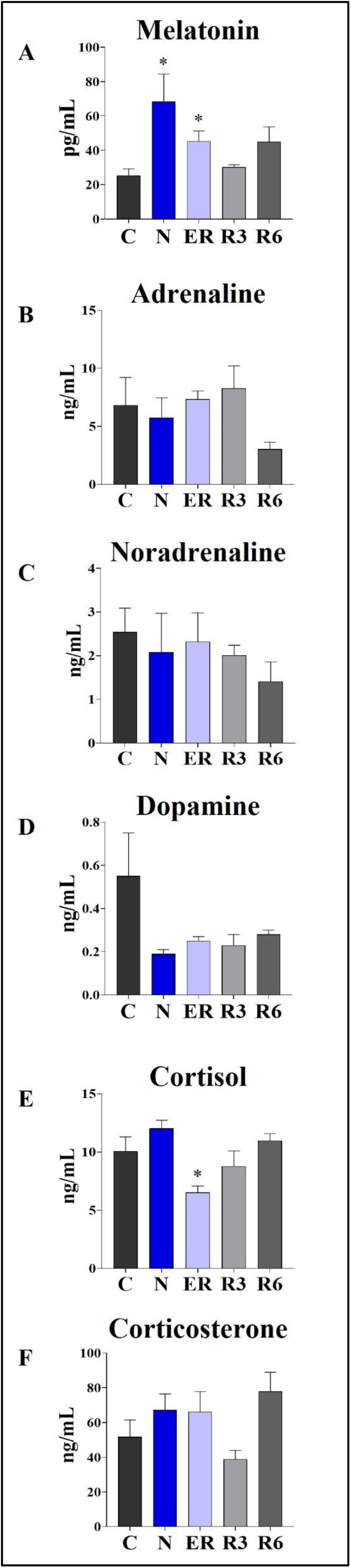
Levels of different plasmatic factors. Panel A: Melatonin; Panel B: Adrenaline; Panel C: Noradrenaline; Panel D: Dopamine; Panel E: Cortisol; Panel F: Corticosterone. Data are expressed as means ± S.E.M., n = 6 (pooled as 3+3, see Methods for details). *: p < 0.05 vs. C. Experimental groups (see Figure 1): C, control; N, samples taken at nadir of hypothermia, during ST; ER, early recovery, samples taken when Tb reached 35.5°C following ST; R3, samples taken 3h after ER; R6, samples taken 6h after ER.

### Microglia morphometry

Figure 7 shows pictures taken from P-Cx during the different experimental conditions (see Fig. 1), as representative samples of the resulting analysis that is summarized in Table 2. Results show that microglia mildly changed some morphological parameters, but only transiently: all the measured values returned to normal within the considered recovery period.

**Table 2.**
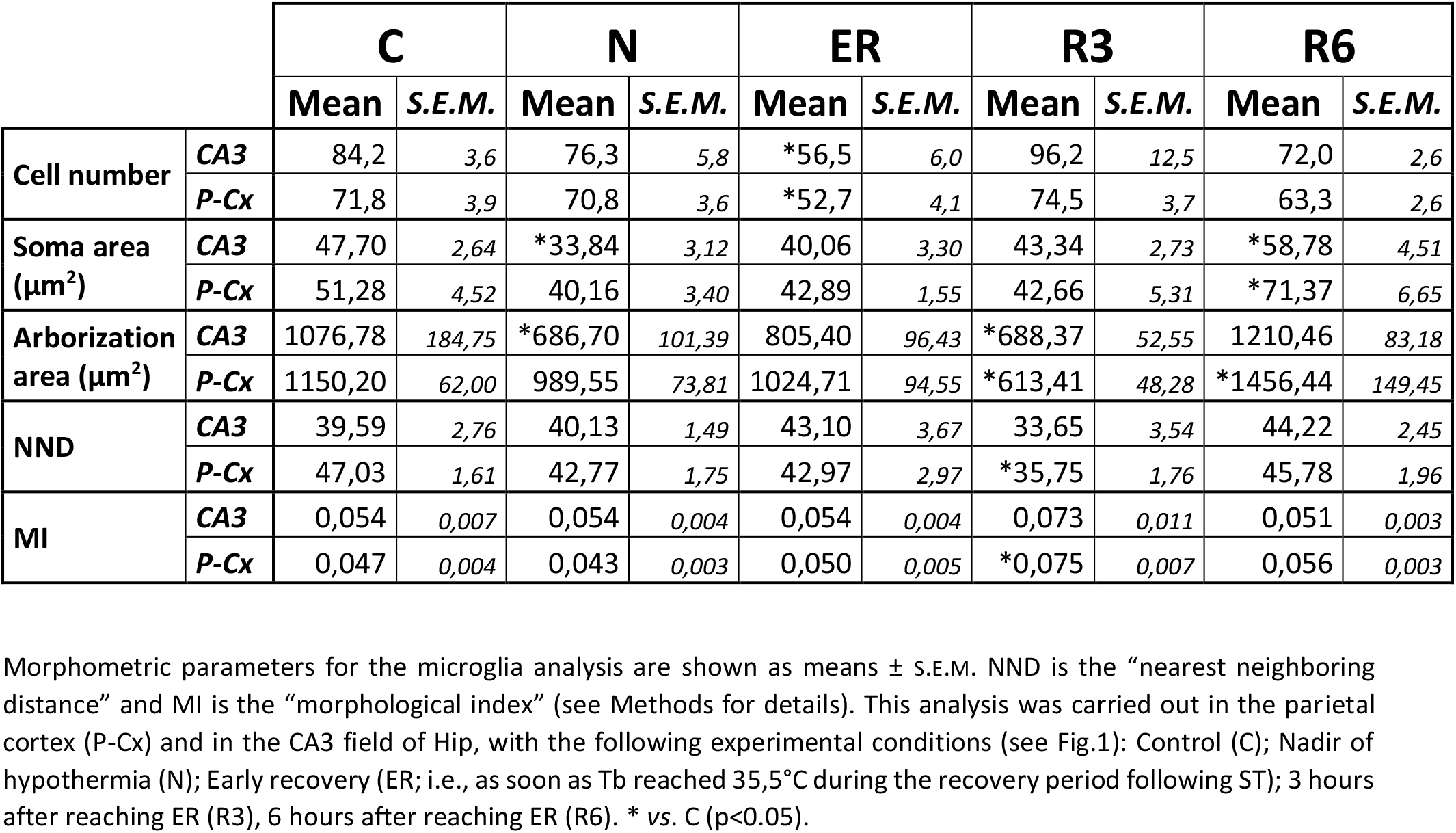
Microglia morphometric analysis

**Figure 7.**
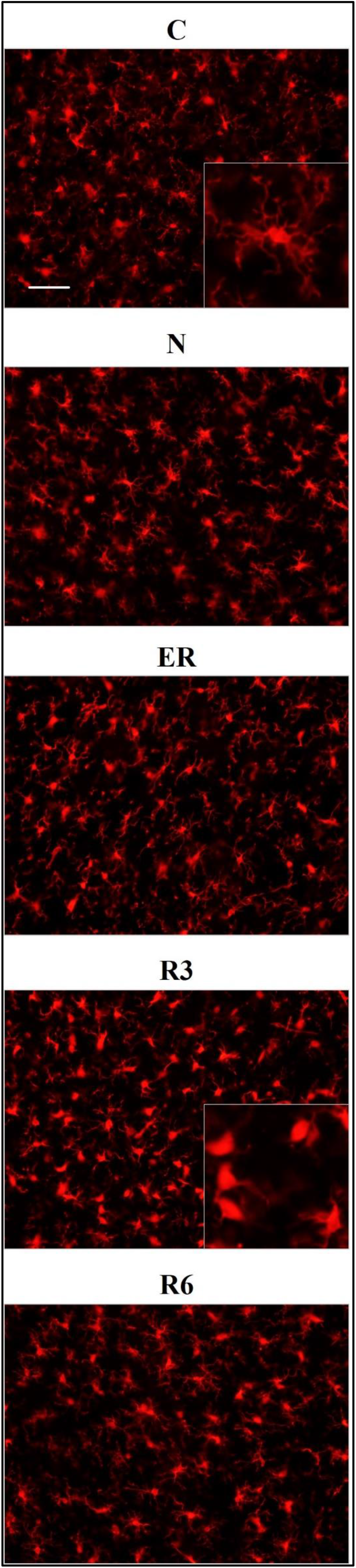
Representative pictures showing microglia, stained for Iba1 (secondary antibody conjugated with Alexa-594), in samples from the parietal cortex (P-Cx) randomly taken between Bregma -2,0 and -4,0. The inclusions within the first (C) and fourth (R3) panels show a ramified and an activated microglial cell, respectively, at higher magnification. Experimental groups (see Figure 1): C, control; N, samples taken at nadir of hypothermia, during ST; ER, early recovery, samples taken when Tb reached 35.5°C following ST; R3, samples taken 3h after ER; R6, samples taken 6h after ER. Calibration bar: 50µm.

In particular, the deep hypothermia reached during ST induced smaller soma and arborization areas compared to C (P<0,05; for both parameters) for microglia in the CA3 field of the Hip (Tab. 2). No differences were found in P-Cx. At the beginning of the recovery period (ER) data showed a lower number of cells compared to C, in both the brain structures studied (P<0,05 for CA3 and P<0,001 for P-Cx). At R3, the arborization area of microglia cells was significantly reduced both in CA3 and P-Cx areas (P<0,05 and P<0,001, respectively), and the NND parameter (i.e., the mean distance between neighboring cells) was lower compared to C (P<0,001), but only in P-Cx. In accordance with these results, at R3 the MI (morphological index) was also found to be significantly higher in P-Cx (P<0,001) compared to C, and close to statistical significance in CA3 (P=0,055). At the end of the recovery period considered (R6), the soma area was larger than in C in both brain structures (P<0,05 for CA3 and P<0,01 for P-Cx), while the arborization area was greater than in C (P<0,05) only in the P-Cx (Tab. 2).

## Discussion

The results of the present work confirm that the reversibility of Tau hyperphosphorylation is not the mere effect of a temperature drop, acting on the physical chemistry characteristics of the enzymatic activity. In fact, the lowering of temperature seems to act as a trigger to elicit an active and regulated biochemical mechanism. Differently from what we supposed, this mechanism does not start to occur during the recovery from ST but it appears to be already strongly activated at the nadir of hypothermia (i.e., the N condition), when Tb is close to 22 °C (Cerri et al., 2013). This was unexpected since rats are non-hibernating mammals, and it was not obvious they evolved some biochemical mechanisms that may act at the low Tb reached during ST. Interestingly, the increase in melatonin plasma levels parallels the changes in the regulatory processes of the enzymes responsible for PPTau dephosphorylation observed during ST and in the following recovery of normothermia, suggesting the possible involvement of melatonin in this neuroprotective process. Another unexpected result was that catecholamine plasma levels were not affected by the experimental procedure, since the returning to euthermia is characterized by thermogenic activation (Cerri et al., 2013; Morrison & Nakamura, 2019). Regarding neuroinflammation, ST induced a mild and transitory activation of microglia cells, indicating their possible role in the recovery of normal conditions of Tau protein.

The present results reflect very well what had previously been observed in our lab in terms of IF determinations (Luppi et al., 2019). In particular, a strong accumulation of AT8 (i.e., p[S202/T205]-Tau) was mirrored by reduced levels of Tau-1 (i.e., the non-phosphorylated form of Tau) in the N condition, with both recovering control levels within the following 3 hours. Moreover, during the recovery period, high levels of Total-Tau were observed, particularly in the P-Cx, indicating a possible stimulating effect of ST on the synthesis of new Tau monomers. The consequences of an enhanced expression of Tau are not easily predictable, since it may be considered a factor that favors either neurotoxicity or cellular neuroprotection (Esclaire et al., 1997; Lesort et al., 1997; Joseph et al., 2017). We believe that, at least in our experimental conditions, the latter is more likely the case. As a matter of fact, p[T205]-Tau form was also shown to be increased at N, also persisting at ER before returning to control levels: this specific phosphorylated form of Tau protein has been described as having a neuroprotective effect in relation to the neurotoxicity induced by β-amyloid in an animal model of AD (Ittner et al., 2016).

Even though the concomitant high peaks of AT8 and p[T205]-Tau in the N condition could be explained, at least in part, by the occurrence of a cross reaction of the two primary antibodies with the two antigens (i.e., AT8 recognizes Tau only when phosphorylated at both S202 and T205 [Malia et al., 2016]), the persistence of high levels of p[T205]-Tau at ER represent a sign of a specific ongoing process with neuroprotective effects. This is supported by the peculiar p[T205]-Tau immunostaining of the CA3 field of the Hip, observed during the N condition, with no staining in the CA1 and CA2 fields (see Figure 3). This result is consistent with the different involvement of the various Hip cortical fields described in AD neurodegeneration (Padurariu et al., 2012; Ugolini et al., 2018). Therefore, the possible neuroprotective process triggered by ST within the Hip is apparently region-specific.

In the present work, we decided to focus on the most representative biochemical pathways involved in Tau phospho-regulation (Planel et al., 2004; Su et al., 2008). Overall, the results from the molecular quantifications of GSK3β and PP2A show that ST elicits a temporary neuronal formation of PPTau that is finely regulated at a biochemical level, as already described in hibernators (Su et al., 2008) and mice (Okawa et al., 2003; Planel et al., 2004; Planel et al., 2007). However, it is worth noting that mice are able to enter torpor as well (Hudson & Scott, 1979; Oelkrug et al., 2011; Hitrec et al., 2019). When planning the experiments, our hypothesis was that some regulated cellular mechanism, that is able to cope with Tau hyperphosphorylation, would take place only during the recovery period from ST, since rats do not hibernate and they might have not evolved specific adaptive mechanisms to cope with torpor bouts (Planel et al., 2004; Su et al., 2008). The present results show that even in the N condition, concomitantly with the excessive PPTau formation, there is a massive biochemical inhibition of GSK3β through phosphorylation on S9 residue (Cross et al., 1995). This response was not expected in a non-hibernator, considering that the enzymatic activity is generally depressed at low temperatures (Aloia & Raison, 1989; Marshall, 1997), with the exception of the specifically adaptive modifications in some enzymatic activity described in hibernating animals (Aloia & Raison, 1989; Su et al., 2008). Moreover, the inhibition of GSK3β apparently persisted, evidenced by the tendency to also maintain high levels of p[S9]-GSK3β during ER in both P-Cx and Hip (Fig. 4). The ER condition also showed a higher PP2A level in P-Cx, a condition that possibly favors the ongoing PPTau dephosphorylation. Hence, this process appeared to be well regulated and important for the quick recovery of normal Tau phosphorylation levels.

In order to verify whether the mechanism elicited by ST is effectively neuroprotective or potentially neurotoxic, we also quantified some key molecular factors involved in apoptosis (i.e., factors that either stimulate or contrast it) or stressogenic cellular processes. Notably, apoptosis represents the main process that causes neurodegeneration (Fricker et al., 2018). In general, changes related to these factors were mild and transitory, such as, for instance, the peak value shown for GRP78 at ER in Hip. However, it is worth noting that when significant differences compared to control levels were observed in these molecular parameters, they were in a direction which indicated a possible neuroprotective rather than a neurotoxic effect. This trend was also confirmed by the observed decrease in the IF staining of cleaved-Caspase 3 in R3, shown in Figure S1. Along the same lines we should consider p[S473]-Akt (i.e., the activated and protective form of Akt [Risso et al., 2015; Levenga et al., 2017]) at ER in Hip.

Considering the specific patterns of data observed in the two brain structures, some interesting differences emerged between P-Cx and Hip: Tau-1 returned to normal values later in Hip than in P-Cx, while GRP78 and p[S473]-Akt only peaked in Hip, for instance. These patterns, on the whole, show that Hip seems to regain normal conditions after ST with more difficulty than P-Cx. This is also corroborated by the quantifications that were observed in AT8 levels; they were much higher in Hip than in P-Cx. Higher values of staining intensity in the Hip, compared to P-Cx, were also found in our previous study using IF (Luppi et al., 2019). Therefore, ST appears to be more stressful to Hip compared to P-Cx, and this is in line with the normally observed evolution of tauopathy within the brain (Crary et al., 2014; Busche & Hyman 2020). However, the difference between the two structures studied in regaining normality was limited to the first few hours of the recovery period; at R6 all the molecular parameters considered returned to normal in both brain areas.

The aim of the present work was also to look for possible systemic factors implicated in the neuronal formation and resolution of PPTau. We mainly focused on plasma melatonin, due to the involvement of this hormone in natural torpor arousal (Willis & Wilcox, 2014) as well as to its capacity to exert neuroprotective effects (Herrera-Arozamena et al., 2016; Shukla et al., 2017). Plasma catecholamines and corticosteroid levels were also assessed, since they might be affected by the strong sympathetic activation which occurs during the recovery of normothermia following ST (Saaresranta & Polo, 2003; Cerri et al., 2013). However, among all the molecules that were tested, only melatonin was shown to change its plasma levels in relation to the experimental conditions. This strongly suggests a possible involvement of this hormone in the neuroprotective mechanism elicited by ST, although we are aware that further experiments are needed to confirm such a hypothesis. In fact, at N the pineal hormone showed a dramatic increase that was concomitant with the peak of p[S9]-GSK3β, suggesting that melatonin may play a role in the process of phosphorylation/dephosphorylation of Tau observed during and after ST. As a matter of fact, a melatonin-mediated neuroprotective effect, leading to GSK3β inhibition and Akt activation, has been described (Liu et al., 2015; Chinchalongporn et al., 2018). The most peculiar aspect of this process, as shown by the present results, is that it is not only activated during the recovery period from ST, as initially supposed, but it starts in deep hypothermic conditions. Since melatonin by itself did not turn out to be effective in contrasting tauopathies (Sanchez-Barcelo et al., 2017), this may suggest that, at a low body temperature, the pineal hormone could act differently from how it acts during euthermia. Indeed, melatonin mainly acts on neurons by means of specific membrane receptors (Liu et al., 2015), but a small quota may directly cross the cell membrane and interact with different cytoplasmatic targets (Liu et al., 2019). A possible explanation of how a low temperature may emphasize the neuroprotective effect of melatonin is the induction of a functional differentiation between these two action modalities. This possibility is supported by data from Chong and Sugden (1994), who studied the thermodynamics of melatonin-receptor binding processes and found that the best affinity ligand-target is actually reached at 21°C, corresponding with the Tb reached by rats during ST in the N condition. Moreover, these authors also reason that at this temperature plasma membrane is very close to a phase transition (Chong & Sugden, 1994). It follows that, at deep hypothermic condition, melatonin could not cross the cell membrane with the same efficiency shown at euthermia. Hence, the coexistence of a much higher binding affinity with membrane receptors (that trigger neuroprotective/anti-apoptotic molecular pathways; Liu et al., 2015), and the difficulty in crossing the cell membrane, may lead to a functional differentiation that could take place only close to 21°C, and not at 37°C. Moreover, the possible involvement of melatonin as a factor in the neuroprotective mechanism elicited by ST is also corroborated by the specific immunostaining of the CA3 field of Hip for the neuroprotective p[T205]-Tau (Ittner et al., 2016) that we observed in the N condition (see Figure 3): in fact, a different regional distribution of melatonin receptors has been described in the Hip, with a higher density exactly in the CA3 field (Lacoste et al., 2015).

Neuroinflammation represents a key condition in tauopathies (Ransohoff, 2016; Nilson et al., 2017), and is also a possible mechanism that induces, or emphasizes, neurodegeneration (Yu et al., 2021). The relationship between neuroinflammation and neurodegenerative pathology is complex: even though neuroinflammation is linked to amyloidosis and tauopathy, it is not clear whether one triggers the other or vice-versa (Guerrero et al., 2021). Apparently, neuroinflammation may be protective as an initial response to a newly developing neuropathology, but, undoubtedly, a sustained chronic inflammatory response contributes to the evolution of neurological diseases (Guerrero et al., 2021). Also, at least in a condition of overt tauopathy, microglia cells appear to actively participate in spreading Tau aggregates (Asai et al., 2015). Our results show that ST induces a mild and temporary microglia activation, mainly shown by the higher MI in Hip and smaller arborization areas observed in both P-Cx and Hip in R3. In a previous work, we also observed a transient microgliosis at R6, that returned to normal after 38h of recovery (Luppi et al., 2019), not confirmed here. Nevertheless, taken together, these results confirm that microglia cells are involved in the resolution of PPTau brain formation during recovery from ST, but microglia activation does not last more than a few hours, since PPTau disappeared. Hence, ST does not trigger a pathological neuroinflammation, but, rather, triggers an acute response that has a possible protective role (Guerrero et al., 2021). This further strengthens the parallelism between ST and hibernation, since a similar pattern of microglia activation has been described in the Syrian hamster (Cogut et al., 2018).

## Conclusion

Deep hypothermia, in rats, seems to uncover a latent and evolutionary preserved physiological mechanism that is able to counteract brain PPTau formation and the evolvement toward neurotoxic effects, strongly supporting the similarity between ST and natural torpor. Considering that the pharmacological stabilization of MTs to treat AD has recently been suggested (Fernandez-Valenzuela et al., 2020), but also that these drugs (which are mainly used as anti-cancer treatments) are highly toxic (Majcher et al., 2018), the indirect stabilization of MTs, as occurs following ST, might be a suitable therapeutic alternative (Craddock et al., 2012a). Thus, the full understanding of what happens at a molecular level in the regulation of Tau phosphorylation/dephosphorylation in neurons exposed to deep hypothermic conditions may assist in the development of new pharmacological approaches to simulate these processes in euthermic conditions, hopefully opening new avenues for the treatment of tauopathies.

## Non-common abbreviations

aCSF: artificial cerebrospinal fluid
AT8: Tau protein phosphorylated at S202 and T205
C: control group
ER: early recovery
Hip: hippocampus
IF: immunofluorescence
MI: morphological index
MTs: microtubules
N: nadir of hypothermia
NND: nearest neighboring distance
P-Cx: parietal cortex
PPTau: hyperphosphorylated tau protein
R3: 3h after ER
R6: 6h after ER
RPa: Raphe Pallidus
Ta: ambient temperature
Tau-1: Tau protein with no phosphorylation within residues 189-207
Tb: deep brain temperature
WB: western-blot

## Acknowledgements

The authors wish to thank: Ms. Melissa Stott for reviewing the English, and Prof. Michelangelo Fiorentini, director of the Operative Unit of Pathologic Anatomy, Ospedale Maggiore, Bologna (Italy), for making a fluorescence microscope available.

## Author contributions

**FS, TH, EP, AO:** Conducted the experiments, collected data. **CG, AO, LT:** Analyzed data, prepared the figures and tables. **ML:** conducted the statistical analyses. **ML, RA, MC**: Conceptualized the experiments. **ML, RA**: wrote the original draft. **All the authors** contributed to the revision and the editing of the paper.

## Original data availability

All the original data are accessible upon reasonable request to the Corresponding author. All the images from western-blot analysis are available on AMS Acta, the Open Science repository of the University of Bologna (http://amsacta.unibo.it/id/eprint/6884 - DOI: 10.6092/unibo/amsacta/6884). These data are under temporary embargo, until a peer-reviewed version of the paper will be published.

## Supplementary Information

Figure S1

