## Supplemental Figure S1 for "Synthetic torpor triggers a neuroprotective and regulated mechanism in the rat brain, favoring the reversibility of Tau protein hyperphosphorylation"

### Supplementary figure

**Supplementary Figure 1.** Representative pictures showing samples from the parietal cortex (P-Cx), randomly taken between Bregma -2,0 and -4,0, stained for cleaved-Caspase 3 (secondary antibody conjugated with Alexa-594). Experimental groups (see Figure 1): C, control; N, samples taken at nadir of hypothermia, during synthetic torpor (ST); R3, samples taken 3h after returning to euthermia; R6, samples taken 6h after returning to euthermia. Calibration bar: 50 µm.

**C**

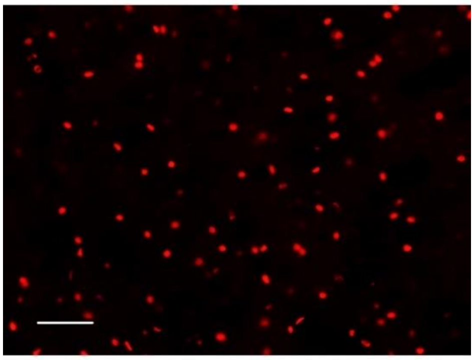

**N**

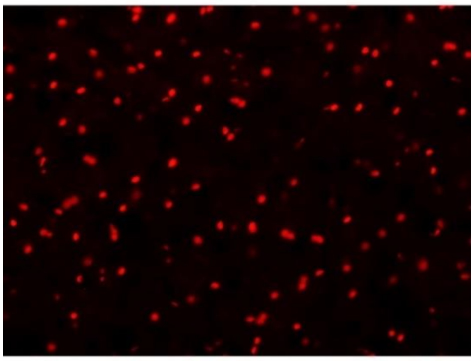

**R3**

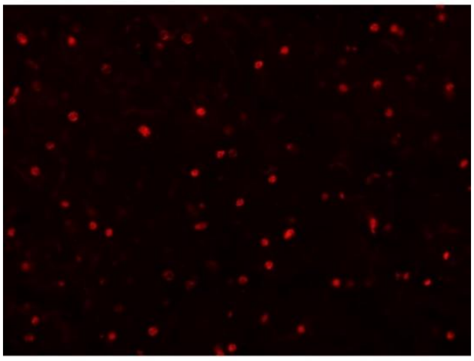

**R6**

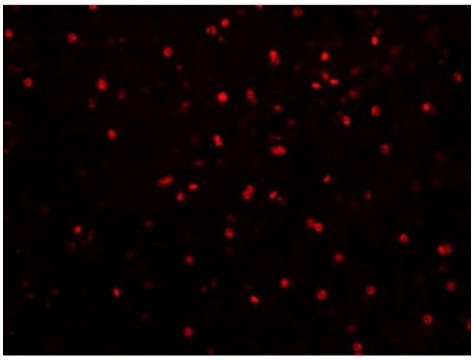
